# Pronouns reactivate conceptual representations in human hippocampal neurons

**DOI:** 10.1101/2024.06.23.600044

**Authors:** D. E. Dijksterhuis, M. W. Self, J. K. Possel, J. C. Peters, E.C.W. van Straaten, S. Idema, J. C. Baaijen, S. M. A. van der Salm, E.J. Aarnoutse, N. C. E. van Klink, P. van Eijsden, S. Hanslmayr, R. Chelvarajah, F. Roux, L. D. Kolibius, V. Sawlani, D. T. Rollings, S. Dehaene, P. R. Roelfsema

## Abstract

During discourse comprehension, every new word adds to an evolving representation of meaning that accumulates over consecutive sentences and constrains the next words. To minimize repetition and utterance length, languages use pronouns, like the word ‘she’, to refer to nouns and phrases that were previously introduced. It has been suggested that language comprehension requires that pronouns activate the same neuronal representations as the nouns themselves. Here, we test this hypothesis by recording from individual neurons in the human hippocampus during a reading task. We found that cells that are selective to a particular noun are later reactivated by pronouns that refer to the cells’ preferred noun. These results imply that concept cells contribute to a rapid and dynamic semantic memory network which is recruited during language comprehension. This study uniquely demonstrates, at the single-cell level, how memory and language are linked.

---

Consider these two sentences: “John and Mary walked into a bar. He sat down at a table.”. When we read the pronoun ‘he’, we realize that John, the only male character in the story, must be the person who sat down at a table. In linguistic terms, John is the ‘antecedent’ of the pronoun. This example illustrates how a narrative activates successive concepts in our brain, including their interrelations, allowing us to incrementally build up a conceptual representation of the discourse^1–3^. Previous brain-imaging studies have gained insight into the brain regions that activate during sentence and discourse comprehension^4–11^. However, the resolution of these non-invasive imaging methods does not suffice to track the neuronal assemblies that encode individual concepts in the human brain during reading.

In recent years, it has become possible to directly record the activity of single neurons in patients who are implanted with electrodes to locate the source of their epilepsy^12,13^. These studies demonstrated the existence of ‘concept cells’ in the medial temporal lobe^14–16^. Concept cells have an invariant and multimodal selective response to a concept. They contribute to the representation of meaning because they not only activate when the participant sees a picture of a specific individual for example, but also when the participant hears or reads the name of this person, or recalls this individual from memory^14,17–19^. We hypothesized that monitoring the activity of concept cells during reading could provide insight into the dynamics of semantic representations during language comprehension. In the present study, we illustrate the possibilities of this approach by examining how pronouns that are encountered during reading influence the neuronal representation of concepts that were introduced in an earlier sentence. Specifically, we asked if pronouns influence the activity of hippocampal neurons. Our results reveal that hippocampal cells respond robustly to preferred nouns and that they are specifically reactivated by a pronoun that refers to them as antecedent. The results represent the first measurements of the magnitude, latency and duration of hippocampal single cell responses to nouns and pronouns during reading, while participants incrementally build up a semantic representation of a narrative.

We recorded from patients with pharmacologically intractable epilepsy who were implanted with depth electrodes in the hippocampus, to localize seizure foci for clinical purposes. During an initial screening session, the patients viewed many pictures of celebrities and family members, and we identified pictures that, for a given neuron, elicited a significantly higher response than other pictures (Methods). We used the results of the screening session, aiming to select three nouns for the main reading task: one was the preferred person for a cell (the preferred noun) and two nouns referred to a male and a female that did not activate that cell (although the non-preferred nouns sometimes activated other, simultaneously recorded cells).

In the main task, on every trial, patients read two sentences which were presented as a stream of words on a computer screen (Fig. 1A and Table 1). The subjects of the first sentence were always two of the three selected nouns (e.g. Courtney Love and Barack Obama), and the second sentence started with a pronoun (either she or he). We then asked a question to verify that the patients understood the meaning of the sentences and maintained focus throughout the experiment. The mean accuracy was 89 ± 2% (s.e.m), significantly higher that the chance level of 33% (Wilcoxon signed rank test, *p* = 1.2·10^-4^; Methods and Fig. 1B).

**Fig. 1.**
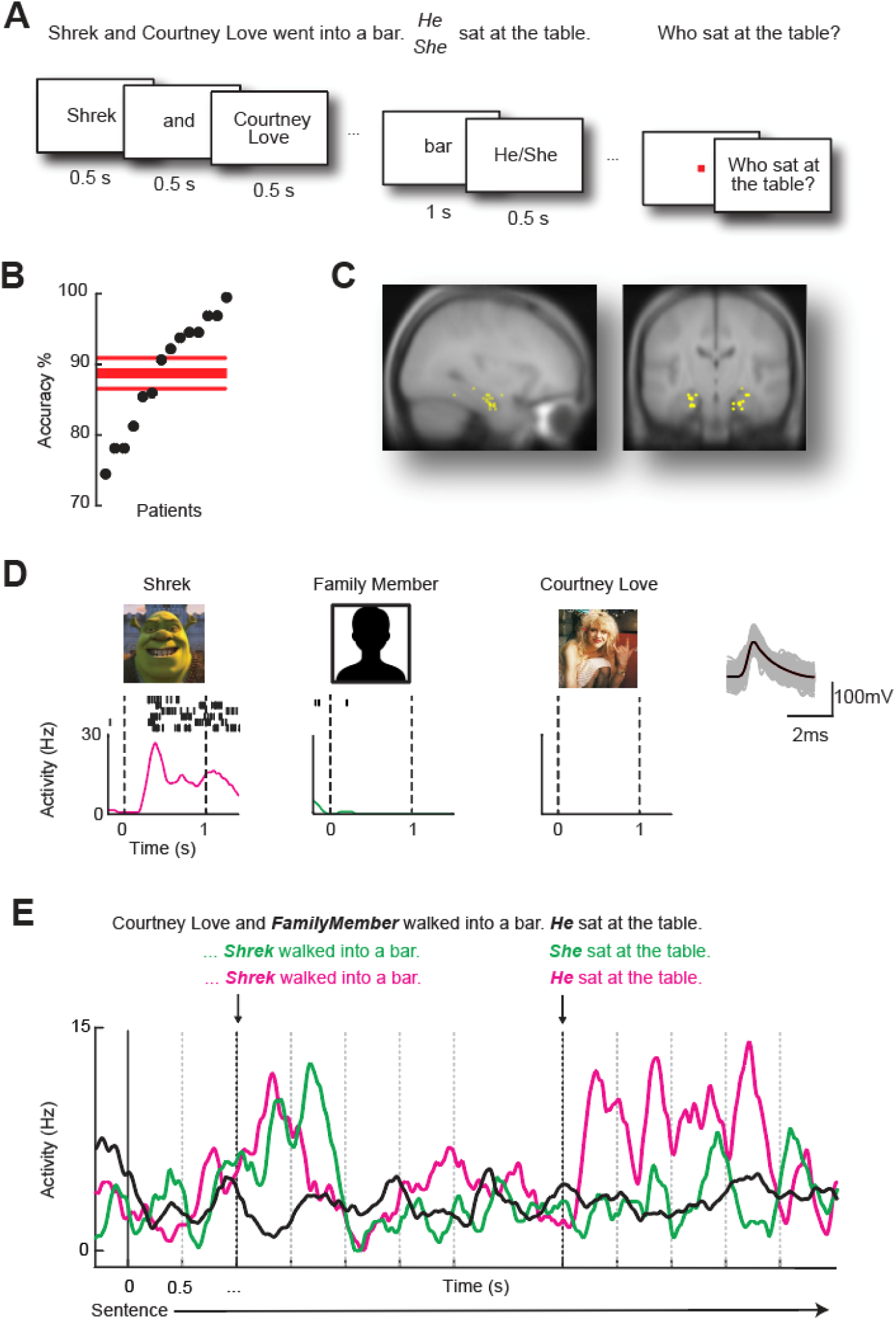
Reading task. **(A)** Two sentences were presented word by word. The first sentence contained two nouns representing individuals (Table 1 and Table S1). The second sentence started with a pronoun. The trial ended with a question (see Methods). **(B)** Accuracy in answering the question across all patients. Red lines, accuracy ± s.e.m. (chance level was 33%, *N* = 14). **(C)** Locations of the electrodes in the hippocampus. **(D)** The average response of an example neuron to different pictures during the screening session, and its waveshape. **(E)** The average response of the ‘Shrek’ cell during the reading task. Sentences in which Shrek appeared as the second noun were followed by a second sentence with the pronoun ‘he’ (magenta) or ‘she’ (green). Activity on trials in which Shrek did not appear in the first sentence is shown in black. Black line at time zero, start of first sentence. Dashed grey lines, onset of successive words. Black dashed lines, presentation of _Shrek_ (or the family member) and the pronoun. The activity elicited by the pronoun was significantly stronger if it referred to Shrek than if it did not (permutation-based Poisson ANOVA, *P* < 0.001).

**Table 1.** Four different contexts in which the noun and pronouns were used for English patients.

| Contexts | First sentences: | Second sentences: |
| --- | --- | --- |
| English | *Noun<sub>1</sub>* and *noun<sub>2</sub>* walked into the bar | *Pronoun* sat at the table |
|  | *Noun<sub>1</sub>* and *noun<sub>2</sub>* sat in the park | *Pronoun* put some sunglasses on |
|  | *Noun<sub>1</sub>* and *noun<sub>2</sub>* were watching the TV | *Pronoun* suddenly changed the channel |
|  | *Noun<sub>1</sub>* and *noun<sub>2</sub>* were having dinner | *Pronoun* poured out some wine |

We identified a total of 529 single and multi-units in the hippocampus of 22 participants (see Table S3 and Methods). We selected cells for further analyses based on their response during the reading task, in which 307 cells responded to the nouns. We focused our analysis on 53 noun-selective cells (recorded in 14 of the participants), which responded significantly stronger to a preferred noun than to the other two nouns (*p* < 0.05, permutation-based Poisson ANOVA and post-hoc Poisson t-test, see Methods) (Fig. 1C). Of the noun-selective cells, we defined 19 to be concept cells, because they preferred the same concept as a picture in the screening task and as a word during the reading task (Methods, Fig. S1 and Table S2; we note that a previous study included more extensive tests of multimodality^20^). The remaining 34 cells were noun-selective, but either did not respond to pictures or preferred a different concept during the screening task, and we will refer to them as noun-selective, non-concept cells (NSNCs). Figure 1D illustrates the activity of an example concept cell that responded selectively to a picture of the animated male movie character Shrek during the screening task. The cell also preferred the written noun ‘Shrek’ (Noun_Pref_) (Figure 1E), but it did not respond to Courtney Love and a male family member of the participant (Nouns_NonPref_). The pronoun ‘he’ in the second sentence activated the cell if it referred to Shrek in the first sentence (Pronoun_Ref_). The pronoun ‘she’ referred to the other noun (Pronoun_PresNotRef_; Table 2 specifies the naming conventions) and did not activate the cell. Furthermore, ‘he’ also did not activate the cell if Shrek was absent from the first sentence (Pronoun_AbsentSame_). Hence, only pronouns referring to Shrek activated the neuron.

**Table 2.** Noun and pronoun conditions. The trials with Pronoun_Absent_ include all trials with Pronoun_AbsentSame_ and Pronoun_AbsentOppo_. Similarly, the trials with Pronoun_NotRef_ include all trials with Pronoun_Absent_ and Pronoun_PresNotRef_. *In two sessions of one of the patients (a total of three hippocampal neurons) we recorded only half of the number of trials.

| Condition description | Nouns in the first sentence | Pronoun in the second sentence | Number of repeats* |
| --- | --- | --- | --- |
| The first sentence contains the preferred noun and a noun of the opposite gender. The pronoun refers to the preferred noun. | Noun<sub>Pref</sub> + Noun<sub>NonPref</sub> | Pronoun<sub>Ref</sub> | 16 |
| The first sentence contains the preferred noun and a noun of the opposite gender. The pronoun refers to the noun with the opposite gender. | Noun<sub>Pref</sub> + Noun<sub>NonPref</sub> | Pronoun<sub>PresNotRef</sub> | 16 |
| The preferred noun is not present in the first sentence. A noun of the opposite and of the same gender as the preferred noun are present. | Noun<sub>NonPref1</sub> +<br>Noun<sub>NonPref2</sub> | Pronoun<sub>Absent</sub> | 32 |
| - The pronoun refers to the Noun with the same gender |  | Pronoun<sub>AbsentSame</sub> | 16 |
| - The pronoun refers to the noun with the opposite gender |  | Pronoun<sub>AbsentOppo</sub> | 16 |
| All trials where the pronoun does not refer to Noun<sub>Pref</sub>. Noun<sub>Pref</sub> can be absent or present. | Noun<sub>Pref</sub>/Noun<sub>NonPref</sub> | Pronoun<sub>NotRef</sub> | 48 |
| The first sentence contains the preferred noun and a noun of the same gender. The pronoun is ambiguous. | Noun<sub>Pref</sub> +<br>Noun<sub>NonPrefSameGender</sub> | Chose<sub>Pref</sub> or<br>Chose<sub>NotPref</sub> | 16 |

To examine the generality of this effect, we carried out an analysis across the population of 53 noun-selective cells. We first examined the activity elicited by the nouns with a cross-validation approach. We used half of the trials to determine the preferred noun and the other half for the statistics. Across the population, the activity elicited by Noun_Pref_ was 5.4 ± 0.5 Hz (mean ± s.e.m.), which was significantly higher than the activity of 2.9 ± 0.5 Hz elicited by the other, non-preferred nouns (t-test, *t* = 3.9, *p* = 3.0·10^-4^). We determined the neurons’ response latencies as the first of 10 subsequent time-bins where the response to Noun_Pref_ was higher than the response to the other nouns (*P* < 0.05; see Methods and Fig. 2D-F). Regardless of whether Noun_Pref_ was presented at the first or second position in the first sentence, the cells activated on average 210ms ± 38ms (s.e.m.) after the presentation of the preferred noun.

**Fig. 2.**
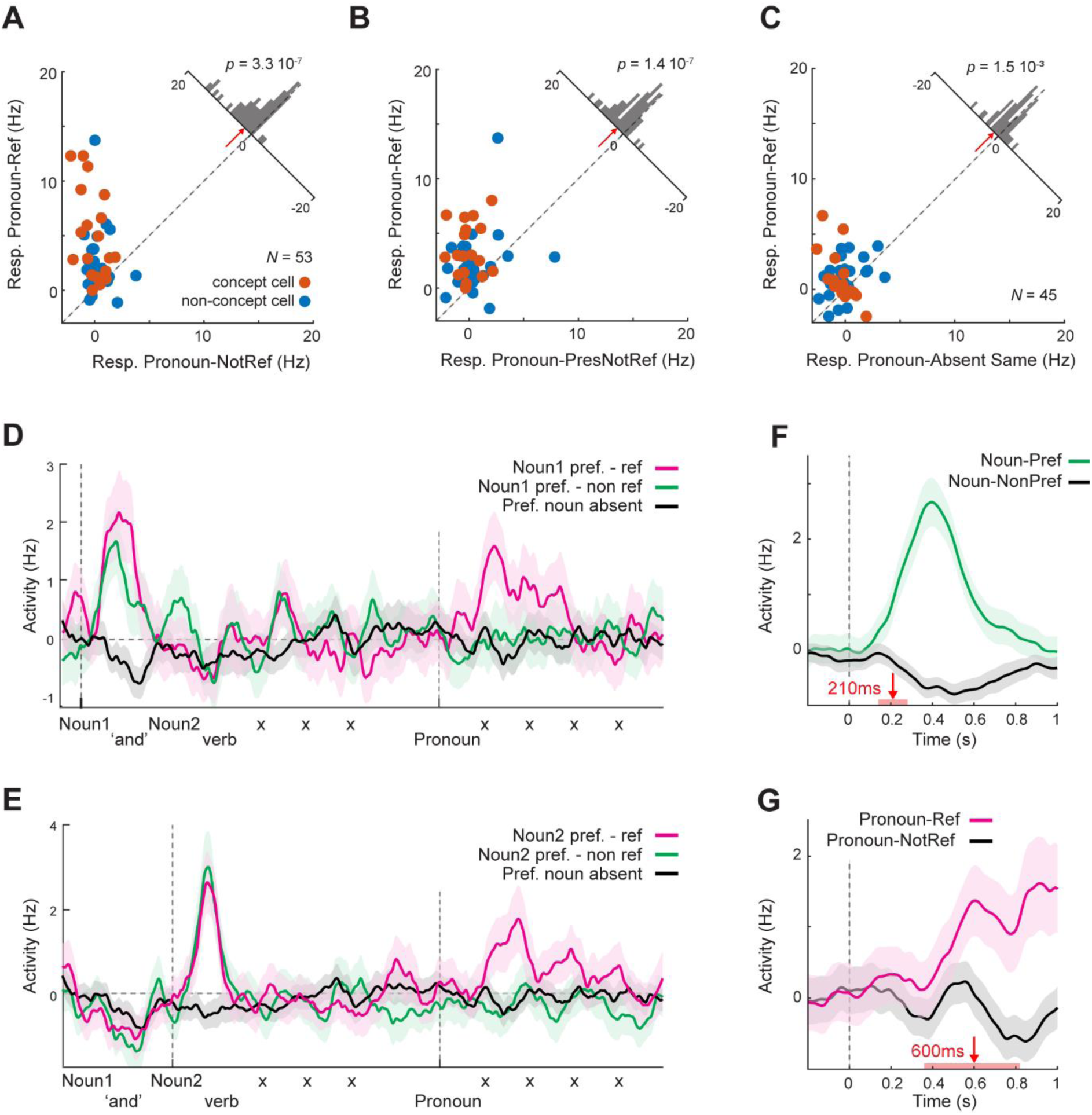
Hippocampal responses elicited by nouns and pronouns. **(A)** Comparison of activity elicited by pronouns referring to the preferred noun (Pronoun_Ref_, y-axis) and all pronouns referring to another noun (Pronoun_NonRef_, x-axis) for concept cells (red dots) and NSNC cells (blue dots). The inset shows the response difference. Red arrow, median. P-value refers to a two-tailed paired t-test (*N* = 53, *t* = 5.85, *p* = 3.3·10^-7^). **(B)** Response to Pronoun_Ref_ (y-axis) and pronouns referring to the other, non-preferred noun on trials in which Noun_Pref_ was present in the first sentence (Pronoun_PresNotRef_, *t* = 6.09, *p* =1.4·10^-7^**) (C)** Response to Pronoun_Ref_ and the same pronoun on trials in which the preferred noun was absent from the first sentence (Pronoun_AbsentSame_, *t* = 3.39, *p* = 1.5·10^-3^). **(D)** Average activity on trials in which Noun_Pref_ was the first noun of the first sentence (Noun1, magenta and green traces) or was absent (black trace), followed by Pronoun_Ref_ (magenta) or Pronoun_PresNotRef_ (green). Shading denotes s.e.m. The vertical dashed lines indicate Noun1 and pronoun onset. **(E)** Average activity on trials in which Noun_Pref_ was the second noun in the first sentence. **(F)** Average activity elicited by Noun_Pref_ and Noun_NonPref_. The red arrows indicate the latency of the response and the red bars the 95%-confidence interval (determined with bootstrapping). **(G)** Average responses elicited by Pronoun_Ref_ and Pronoun_NotRef_.

We next examined the responses elicited by the pronouns. On average, pronouns referring to Noun_Pref_ (Pronoun_Ref_) activated the hippocampal neurons more strongly than pronouns that did not (paired t-test, *t*_52_ = 6.8, *p* = 3.3·10^-7^) (Fig. 2A). This effect was significant for 25% of the noun-selective cells and for 42% of the concept cells (permutation-based Poisson ANOVA; *P* < 0.05). Interestingly, the responses of concept cells to Pronoun_Ref_ (5.0 ± 0.9 Hz*, N* = 19) were stronger than those of the NSNC cells (2.3 ± 0.5 Hz, *N* = 34; t-test corrected for unequal variance, *t* = 2.6, *p* = 0.016; Fig. 2A). The extra activity elicited by Pronoun_Ref_ emerged gradually (Fig. 2G) with a latency of 600ms ± 110ms, which is longer than the latency of the response to Noun_Pref_ (*p* = 0.010, bootstrapping test), although activity built up gradually so that the inclusion of more cells might have shortened the latency estimate. There were no significant differences in response magnitude (independent samples t-test, *t*_51_ = 0.19, *p* = 0.85) or latency (*p* = 0.54; bootstrap test, see Methods) between the left and right hemispheres.

Is the response to Pronoun_Ref_ caused by a prolonged activation by the preferred noun, extending into the second sentence? To examine this question, we selected the first sentences with Noun_Pref_ and compared the activity elicited by pronouns that referred to Noun_Pref_ (Pronoun_Ref_) and to the other noun (Pronoun_PresNotRef_); for example ‘Shrek and Courtney Love’ followed by ‘He’ or ‘She’. The response elicited by Pronoun_Ref_ was higher than that evoked by Pronoun_PresNotRef_ (paired t-test, *p* = 1.4·10^-7^) (Fig. 2B), indicating that it reflects the antecedent of the pronoun and is not a lingering memory trace of Noun_Pref_.

We next investigated whether the pronoun activation reflected tuning to gender rather than to the identity of the antecedent. We compared trials in which the pronoun referred to Noun_Pref_ (e.g. ‘Shrek and Courtney Love’ followed by ‘He’) to trials in which the same pronoun referred to a different noun of the same gender (Pronoun_AbsentSame_, e.g. ‘Barack Obama and Courtney Love’ followed by ‘He’). Pronoun_Ref_ elicited a higher response than Pronoun_AbsentSame_ (*N* = 45, paired t-test, *p* = 1.5·10^-3^; Fig. 2C), implying that the pronoun responses reflected the identity of the antecedent and not the gender.

We examined whether population responses evoked by pronouns resembled those evoked by the nouns and how different cell groups contributed to the population response. To this end, we trained a linear support-vector machine (SVM) to discriminate between responses elicited by Noun_Pref_ and Noun_NonPref_. We constructed 1,000 surrogate-populations by randomly choosing units from the noun-selective cells with replacement (*N* = 50 cells; we excluded three cells from this analysis, Table S3). We trained the classifier on activity in the noun time-window and tested the classifiers on held-back trials in a series of sliding windows covering the two sentences. We split trials based on first sentences in which Noun_Pref_ 1) appeared at the first position, 2) appeared at the second position and 3) was absent. The classifier output represents the fraction of trials in a particular time-window in which the neuronal responses are labelled as elicited by Noun_Pref_ (y-axis in Fig. 3A). As expected, the classifier output increased shortly after the presentation of Noun_Pref_ at both the first and second position, labelling the activity as evoked by Noun_Pref_ in ∼90% of trials (Fig. 3A). Interestingly, when Noun_Pref_ was at the first position, the classifier output increased slightly after the presentation of the second, non-preferred noun (black arrow in Fig. 3A), which may reflect a mental process that relates the two nouns.

**Fig. 3.**
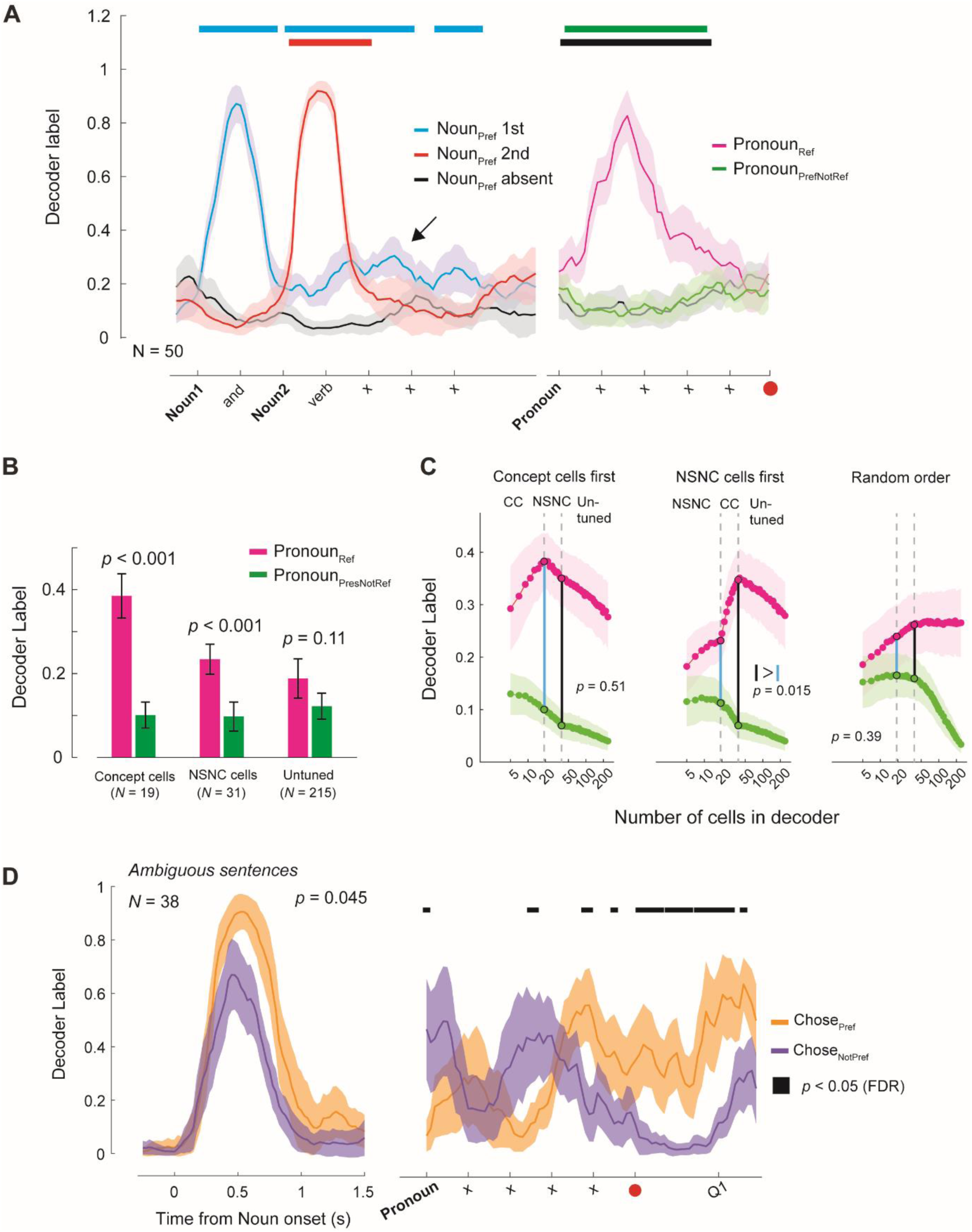
Decoders for the preferred noun. **(A)** SVM classifiers trained to distinguish between the responses to preferred and non-preferred nouns. The output (y-axis) represents the fraction of trials in which the pattern of activity in a time-window is classified as evoked by Noun_Pref_. The classifiers were trained on the activity of 50 cells in the noun time-window and then tested on the two sentences in a series of sliding windows of 0.5s duration. The words are marked at the centre of the sliding window, the red dot indicates the end of the second sentence. The shading around the traces indicates the standard deviation across surrogate populations (*N* = 1,000). Filled bars indicate significant time windows (z-test, *P* < 0.05, FDR corrected; see Methods). **(B)** Decoder output for different cell populations (0.5-1.5 seconds after pronoun onset) for Pronoun_Ref_ (magenta) and Pronoun_PresNotRef_ (green). Error bars, standard deviation across surrogate populations. P-values, z-test on the difference between conditions. **(C)** Classifier output as function of the number of included units. **Left,** We first added concept cells (CC: *N* = 19), then NSNC cells (*N* = 19 cells selected from a total of 31 cells) and then untuned units (*N* = 215). Decoder output is shown for Pronoun_Ref_ (magenta) and Pronoun_PresNotRef_ (green). **Middle,** We first added 19 NSNC cells, then 19 CC cells and then the untuned units. **Right,** We added units in a random order. We compared the decoder output between decoders with only CCs, only NSNCs (*N* = 19, length of blue vertical lines) to that when cells of the other category were added (*N* = 38, length of black lines) with a z-test to determine significance (P-values in the panels). **(D)** Decoder output on ambiguous trials. The decoders, which had been trained on non-ambiguous trials to discriminate between preferred and non-preferred nouns, were tested on ambiguous trials with two nouns of the same gender. The left panel shows the decoder output to the preferred noun in the first sentence (aligned on Noun_Pref_ onset). The right panel shows the decoder output during the second sentence, in sliding windows with a duration of 0.5s. We sorted the trials based on whether the participant selected the preferred noun (Chose_Pref_, orange) or the non-preferred noun of the same gender (Chose_NotPref_, purple) in response to the question. Q1, onset of the question.

We next tested the classifier on the second sentence, on which it had not been trained. We compared the classifier output between trials in which the pronoun referred to the preferred noun (Pronoun_Ref_), trials on which it referred to the other noun (Pronoun_PresNotRef_) and trials on which Noun_Pref_ was absent from the first sentence. The activity pattern elicited by Pronoun_Ref_ was like that elicited by Noun_Pref_, causing a higher decoder output than on trials in which the pronoun referred to the other, non-preferred noun and on trials without Noun_Pref_ in the first sentence (Fig. 3A). The decoder output was also higher than on trials in which the antecedent was the non-preferred noun with the same gender (Pronoun_AbsentSame_; Fig. S2A). We replicated the results with a cross-decoding analysis, training and testing on all combinations of time-bins in the two sentences. Cross-decoding worked in both directions, from noun to pronoun and from pronoun to noun (Fig. S2B).

To investigate which cells contributed most to the selectivity of the pronoun response, we built decoders by successively adding concept cells (*N* = 19), NSNC cells (*N* = 19 selected of a total of 31 units) and neurons that responded to nouns without significant tuning in different orders (‘untuned’, *N* = 215, noun response > 0.5 Hz above baseline; see Table S3 for inclusion of cells) (Fig. 3B). Decoders built from noun-selective cells (concept and NSNC) to decode nouns also reliably decoded the pronoun of the second sentence, whereas adding untuned cells did not improve this form of generalization (Fig. 3C). Interestingly, adding NSNC cells to decoders that already included all concept-cells did not improve decoding, whereas adding concept-cells to decoders with only NSNC cells improved decoding accuracy (*p* = 0.015, z-test) (Fig. 3B,C). These results indicate that decoding of the pronoun’s antecedent relied mostly on the concept cells, with a smaller contribution from NSNC cells and little contribution from the untuned cells.

Two additional observations demonstrated that the neural responses were linked to the subjects’ interpretation of the pronoun on individual trials. First, we compared the response to pronouns that referred to the preferred noun (Pronoun_Ref_) between correct trials and error trials, on which the participants reported the incorrect noun in response to the question at the end of the trial. The hippocampal response was higher on correct trials than on error trials (t-test, two-tailed, *t* = -2.31, *p* = 0.008, *N* = 35; Fig S3A), indicating that it predicted the participant’s interpretation of the second sentence.

Second, we analysed ambiguous trials with two nouns of the same gender in the first sentence, so that they competed for a status as antecedent of the pronoun (‘Shrek and Barack Obama went into a bar. He sat at the table’, followed by the question ‘Who sat at the table?’) (Fig. 3D; Table 2). We predicted that the strength of the response to Noun_Pref_ in the first sentence might relate to the probability that the participant chooses it as antecedent and reports it in response to the question. In accordance with this prediction, the decoder output for the preferred noun of the first sentence was stronger if the participant chose it in response to the question (Fig. 3D) (*p* = 0.045, z-test). Furthermore, decoder output also ramped up at the end of the second sentence if the participants chose the preferred noun (Fig. 3D). Hence, the activity of hippocampal neurons during reading is related to the strength of the conceptual representations and predicts the resolution of pronoun ambiguity.

The results, taken together, demonstrate a link between the activity of neurons in the human hippocampus and the incremental representation of concepts and their interrelations during sentence reading. Previous studies indicated how language understanding relies on the integration of semantic knowledge, to which the hippocampus contributes^21–26^, with linguistic representations^27–31^. We here provided the first measurements of single unit activity in the hippocampus during a task in which participants build up a semantic representation of a narrative^32^, in which a pronoun brings one of the characters into the foreground of thought. Interestingly, most of the information about the antecedent of the pronoun was carried by concept cells, which responded to a concept irrespective of whether it was presented as picture or noun, indicating that their activity disambiguates otherwise identical pronouns.

Hippocampal neurons responded to both their preferred noun and to pronouns referring to that noun. The hippocampal pronoun response could thereby link new information of the narrative to the appropriate concept. For example, when we read about Shrek that ‘he’ put on sunglasses, we can update Shrek’s representation and predict his future appearance^14,33–35^. In accordance with this view, damage to the hippocampus can lead to impairments in pronoun production and comprehension and patients with hippocampal lesions often fail to retrieve a pronoun’s antecedent^36,37^. Our results support a role for the hippocampus in pronoun resolution, because the pronoun response was weaker on error trials and hippocampal activity predicted the perceived antecedent of the pronoun on ambiguous trials.

The latency of the pronoun response was 600ms, which was longer than the latency of 210ms of the noun response, and most likely reflects processes that link the representation of the pronoun in language areas to the appropriate concept in the hippocampus. The long latency is remarkable because pronouns are thought to increase discourse efficiency, allowing shorter utterances to activate the same concepts as longer noun phrases^1,38^. The present results indicate that this pronoun advantage may only occur for noun phrases that take several hundreds of milliseconds. We note, however, that the latency of the pronoun response in the hippocampus might be shorter if there is only a single possible antecedent.

Language theories propose that pronouns have additional advantages, such as signalling that the subject of the narrative remains the same across sentences^39^. Our results inform these theories^1,39^, including ‘prominence’ theories that propose that pronouns usually refer to the most prominent nouns of a discourse^40^. According to those theories, nouns that fulfil a similar role in previous sentences compete for selection based on their prominence, which reflects on how important they are in the narrative^40^. So far, these theories could only be tested indirectly^41,42^, short of a method to measure prominence in the brain. Our results on trials with two nouns of the same gender, which caused the pronoun to be ambiguous, provided insight into the neuronal underpinnings of prominence. On these ambiguous trials, the participant could report either noun as antecedent of the pronoun. When the participants reported that the pronoun referred to the preferred noun, the response to that noun was stronger during the first sentence, and hippocampal activity ramped up once more at the end of the second sentence. These results suggest that prominence is related to the strength of the conceptual representations, which depend, in part, on the activity of hippocampal neurons.

Theories about the evolving mental representation of the narrative during reading suggest that previously read words are stored in working memory so that they can be combined with new information^28–30,32,43,44^. Interestingly, the hippocampal activity elicited by nouns lasted only 300-400ms and was curtailed by later words, which has implications for the internal structure of these representations^28^. Theories of working memory propose that memorized items are not equivalent, because only one or a few of them can be in the focus of attention^45^. Studies in the frontal cortex of monkeys revealed that attended and non-attended items in working memory are represented as sustained activity patterns^46,47^, whereas the present results suggest that the hippocampus selectively represents the attended noun. The activity of the neurons largely vanished during the reading of subsequent words, but the cells became active again in response to a pronoun referring to the cell’s preferred concept. Hypothetically, the representation could have moved into a different neuronal subspace, as observed during sequence working memory in monkey prefrontal cortex^46^, but this is unlikely given that there were epochs in which we could no longer decode it from the population of neurons. The activity profile rather suggests that hippocampal neurons predominantly represent concepts which come in the focus of attention as a central topic of the narrative. These attended concepts could then be used for the retrieval of additional associations, a process to which the hippocampus contributes^48^,

In our experiment, pronoun disambiguation was based on gender alone, but in other sentence constructions, syntax plays an essential role. For instance, in the sentence “The fact that he is so rich pleases John”, the pronoun “he” can refer to John, but in “He is so rich that John is pleased”, it cannot. How brain networks implement such syntactic computations is a topic for future research, which can now be investigated. The present results show how single unit recordings in the medial temporal lobe contribute to the flexible linkage of words and concepts during language comprehension.

## Acknowledgments

We are grateful to the staff of the EMU of the VUmc and UMCU for their help during these recordings. We thank Peterjan Ris and Ndedi Sijsma from the VUmc, and Luigi Rizzi, Naama Friedmann, Hilda Koopman and Klaus von Heusinger for discussions at various stages of this project. We thank Peter Hagoort for his valuable feedback on the manuscript.

## Author contributions

Conceptualization: MWS, PRR, SD

Clinical procedures: JCB, SI, PE, RC, VS, IES, SMAS, EA, NCEK, DTR

Ethical approval acquisition: MWS, DED, SH

Investigation: DED, MWS, JKP, JCP, SH, FR, LK

Formal analysis: DED, MWS

Visualization: DED, MWS

Funding acquisition: MWS, PRR

Supervision: MWS, PRR

Writing – original draft: DED, MWS, PRR

Writing – review & editing: DED, MWS, PRR, SD

Competing interest declarations

None.

## Data availability

All source data and code to create the main and supplementary data figures will become publicly available via EBRAINS (https://ebrains.eu/service/share-data/).

## Supplementary Materials

Methods

Figs. S1 to S5

Tables S1 to S3

