## Supplementary Materials for "Pronouns reactivate conceptual representations in human hippocampal neurons"

**This PDF file includes:**

Methods

Figs. S1 to S3

Tables S1 to S2

### Methods

Participants and electrodes

All 22 participants were patients with pharmacologically intractable epilepsy from the University Medical Centre VU Amsterdam, the University Medical Centre Utrecht (both in the Netherlands) and the Queen Elizabeth Hospital, Birmingham (UK). They were implanted with depth electrodes for chronic seizure recording to examine surgical resection possibilities[12], for 7-14 days. S.I., J.C.B., P.E. and R.C. performed the surgeries. The Medical Ethics Committee at the UMC VU Amsterdam approved the experiments in the Netherlands; the National Health Service Health Research Authority 175 (15/WM/0219) approved the experiments in the UK. Before participation, all patients gave their written informed consent. Solely clinical criteria determined the location of the electrodes, with the aim to determine a resection plan with highest probability of decreasing seizure frequency. The researchers indicated which of the electrodes would be of the Behnke-Fried type, with nine microwires extending at the end of the clinical electrode[12]. Eight of the wires had a high impedance to record single- and multi-unit activity and one served as a low-impedance local reference. The 22 patients were implanted with a total of 49 Behnke-Fried electrodes in the hippocampus, which is the area we focus on in the current study because the number of units in other structures of the medial temporal lobe was too small for our analysis. We confirmed that the microwires were in the hippocampus based on the post-implantation CT scan, in which the macro-contacts were visible, co-registered to the pre-implantation T1-weighted structural MRI scan. We verified for each electrode and participant that the electrode tips fell within the hippocampus (see Fig. S5 for an example patient). For visualization of the electrode locations, we registered the CT and MRI scan to MNI space using SPM12 and placed a 3-mm-radius sphere 3 mm distal to the tip of the macro-electrode (Fig. 1C).

Screening experiment

The first task for the participants was a screening experiment. We presented many pictures (median 86 pictures) of relatives and friends and famous people on a laptop computer, while the participant sat in the hospital room. Each picture was presented at the centre of the screen for 1000ms, occupying 1.5° of visual angle. The inter-stimulus interval (ISI) was 500ms and each image was shown 8 times in a randomized order. We ensured that the participant maintained attention on the pictures by asking a question about the person in the picture, for example, ‘Is the person on the picture Dutch or not Dutch?’ or ‘Is the person on the picture a family member or not?'. The patients could respond with a key press during the presentation of the picture and the subsequent ISI period. The next picture presentation would automatically start independently of the response time of the participant. No feedback was given during the task.

Reading experiment

After the screening session, the patients performed a reading session in their native language. On each trial, two sentences were presented on a black background, word by word (with the exception of certain names, e.g. "Courtney Love" composed of two words, which were presented simultaneously) for 500ms per word (in white) (Fig. 1A). One participant needed more time for reading and the stimulus duration was extended to 600ms. The first sentence contained two proper nouns representing two different individuals and 5 other words (for example: "*Shrek* and *Courtney Love* walked into a bar."). We selected the proper nouns based on the results from the screening session. After an initial analysis of the screening task data and a visual inspection of the patterns of spiking activity we selected three pictures that were most likely to drive selective responses to nouns in the reading task. During this selection process we aimed to maximize the number of responding units while minimizing the chance that a unit would respond to more than one noun. We selected two nouns of one gender and one noun of the opposite gender. Nouns occurred equally often at the first or the third position in the first sentence, referred to as 'noun position 1' and 'noun position 2'. The word in position 4 was always a verb. The last word of the first sentence was presented for 1000ms. The second sentence started with a pronoun, referring to one of the nouns (for example: "*He* sat at the table."). In some trials, two nouns of the same gender appeared in the first sentence. In these cases, the pronoun could refer to either noun and these trials were therefore called ambiguous. After sentence two, a small red fixation dot (0.23°-0.34°, depending on the distance between the participant and the screen) was presented for 1000ms, followed by a question that was displayed on the screen (for example: "Who sat at the table?"; Fig. 1A). After 500ms the three nouns appeared on the screen, with locations that varied across trials. The patient chose one noun by pressing the left-, middle- or right arrow key of a keyboard. They had unlimited time to respond. The average performance accuracy of the patients was compared against chance level (33.3%) with a Wilcoxon signed rank test ($P < 0.05$). Table 1 illustrates the sentences that were used in our study and Table S1 the Dutch versions. There was a blank screen of 500ms after the response of the patient, and then the next trial started. The recording sessions lasted 8-15 minutes with 40 to 80 sentence pairs (see Table 2 for the number of trials per condition).

Spike detection and responsiveness criteria

In total, we recorded from 392 micro-wire electrodes located in the hippocampus during 49 sessions. The signal from the microwires was amplified using impedance-converting head-stages placed on the head of the patient (Neuralynx 'HS-9'/Blackrock 'Cabrio') and it was either recorded with a 64-channel Neuralynx ATLAS system (32 kHz sampling rate) or with a 128-channel Blackrock NeuroPort Biopotential Signal Processing System (30 kHz sampling rate). Digital filters (Neuralynx: high-pass 0.1-1 Hz, low-pass 9000 Hz, Blackrock: high-pass 0.3, low-pass 7500 Hz) were applied after sampling. We used a semi-automated algorithm to determine whether to digitally re-reference the raw signal to a micro-wire from the same bundle. The algorithm attempted to minimise RMS noise levels while also avoiding re-referencing to a micro-wire that exhibited spiking. The choice of reference was made by the experimenter (D.D.). After re-referencing, the data from the screening and corresponding reading session were concatenated and spikes were detected and sorted using semi-automatic methods, as described previously[17,49]. In short, the raw-data was band pass filtered between 300 and 1500 Hz, and an automatic amplitude threshold was applied to detect threshold crossings (usually ~6 times the median absolute deviation of the filtered data time-series). The raw data was then re-filtered between 500-3000 Hz and spike waveforms were extracted around the threshold crossings. The spike waveforms were clustered using a wavelet transform and a previously described algorithm using WaveClus 3[50]. Clusters were visually inspected by D.D. who merged, split or excluded clusters depending on their waveform, signal-to-noise ratio and ISI: single ($N$ = 171) and multi-units ($N$ = 358) in the hippocampus were included in the analysis. We evaluated the quality of isolation of single units by computing the number of inter-spike intervals (ISIs) smaller than 3ms (ISI-violations), which was 0.34% (compared to 1.7% for multi-units) and the SNR, which was 16.6 on average (7.7 for multi-units).

We visualized the neuronal responses with peristimulus time histograms, binning the spike-times (1ms bin width) and convolving them with a Gaussian kernel with a s.d. of 1.5ms. The PSTHs were baseline corrected by subtracting the overall average firing rate over the entire experimental session. We smoothed the average PSTHs with a sliding window of 20ms (MATLAB 'smooth' function).

Analysis of noun and pronoun responses

Data analyses were performed on spike-counts from only correct trials, except where indicated. To determine whether units were noun-selective, we examined whether their response to the three nouns differed significantly with a permutation test based on a Poisson generalized linear model (GLM), as described previously[34]. In brief, we calculated the mean spike count in 15 analysis windows, starting at 200ms after noun onset in steps of 100ms and with durations ranging from 100-600ms, not extending past 800ms. In each window we compared the likelihood of observing the data with different mean firing-rates across nouns ($\mathcal{L}_{Full}$) to the likelihood of the data with the same mean firing rate across nouns ($\mathcal{L}_{Null}$). The likelihoods were derived according to the equations used in Kornblith et al., 2017[34]. We then derived a $\chi^2$ statistic which is equal to the difference between the deviance of a GLM incorporating the effect of stimulus ($D_{full}$) and an intercept-only GLM (i.e. all nouns elicited the same firing rate, $D_{Null}$):

$$\chi^2 = D_{Full} - D_{Null} = 2\log(\mathcal{L}_{Full} - \mathcal{L}_{Null}) \qquad (1)$$

The test statistic was the maximum value of $\chi^2$ across time windows. We also identified the time-window $W_{noun}$, in which the response between the three nouns is maximally different. The test statistic was compared to a null distribution of the maximum $\chi^2$ statistic across time-windows derived from a surrogate dataset made by shuffling the noun labels 1000 times: $\chi^2_{null}$. The p-value was estimated as the proportion of $\max(\chi^2_{null})$ that was greater than or equal to $\chi^2$. We used an alpha level of 0.05 to identify the noun-selective units with significantly different responses across nouns. We applied a post-hoc Poisson-distributed independent samples t-test to determine whether the response to the preferred noun, which gave the maximum response in $W_{noun}$, was significantly greater than the average of the response to the remaining two-nouns. The spike-rate $R_{noun}$ to the preferred noun in $W_{noun}$ is a measure of the response of the unit and we only included units where $R_{noun} > 0.5$ Hz.

We tested noun-selectivity at the population level with a cross-validation approach. We identified the preferred noun of each unit using even-numbered trials and then tested the responses on odd-numbered trials. For each unit we calculated $R_{noun}$ using the odd-numbered trials for both the preferred noun and average across the non-preferred nouns. The difference in this metric across neurons was normally distributed and was tested using a two-tailed paired t-test.

We examined the significance of the activity elicited by pronouns using a similar approach. We determined the spike-rate in 28 analysis windows ranging from 200-1000ms after the onset of $\text{Pronoun}_{\text{Ref}}$ and $\text{Pronoun}_{\text{NotRef}}$. These time windows were longer than for the nouns because we expected *a priori* that the pronoun response might be delayed relative to the noun. We identified the time-window $W_{pronoun}$ with the maximum average response to $\text{Pronoun}_{\text{Ref}}$ and $\text{Pronoun}_{\text{NotRef}}$, an approach that does not bias the population statistic as the two conditions contributed equally to the average response. We used the firing rate in $W_{pronoun}$ elicited by the different pronoun types as the statistic for the population analysis (Figures 2A-C) using two-tailed paired t-tests. To compare the activity elicited by $\text{Pronoun}_{\text{Ref}}$ and $\text{Pronoun}_{\text{AbsentSame}}$, we excluded units whose preferred noun was the noun with the unique gender ($N = 8$) because the $\text{Pronoun}_{\text{AbsentSame}}$ could not be tested in these units. We carried out the Poisson permutation test with 10,000 shuffles to generate p-values for each unit.

To examine whether there was a relation between the activity elicited by the pronoun and the participant's response to the question after the second sentence, we split trials into correct and error trials for the $\text{Pronoun}_{\text{Ref}}$. Cells from sessions with an insufficient number of incorrect trials were excluded. The difference between the response to $\text{Pronoun}_{\text{Ref}}$ on correct trials versus incorrect trials was tested with a two tailed t-test (Fig. S3A).

Description of the population of hippocampal units

Of a total of 529 single and multi-units in the hippocampus, recorded in 22 participants, 307 responded to nouns with $R_{noun} > 0.5$ Hz (Table S2). Most of our analyses focused on the 53 noun-selective cells (40 multi- and 13 single units), recorded in 14 of the participants (26 sessions out of a total of 49 sessions across all participants, Table S3). Of these noun-selective units, 28 came from the left hippocampus and 25 from the right hippocampus. The magnitude of the activity elicited by pronouns referring to the preferred noun was very similar in the two hemispheres, 3.2 Hz and 3.0 Hz for the left and right hippocampus, respectively (independent samples t-test, $t_{51} = 0.19$, $p = 0.85$). There was no difference in response latency between the hemispheres either ($p = 0.54$; bootstrap test). Hence, the effect is present bilaterally without a clear hemispheric asymmetry, in line with previous work[51].

To determine which of noun-selective units were concept cells, we compared selectivity to the nouns during the reading session to selectivity to pictures during the screening session, using the analysis described above for nouns to determine picture selectivity. Cells were considered picture selective if images elicited different response magnitudes ($p < 0.05$, permutation-based Poisson ANOVA), and if the elicited response to at least one and maximally five pictures was stronger than that elicited by the other pictures ($p < 0.05$, post-hoc Poisson t-test). We considered 19 units to be concept cells, because they were selective to a picture of the same person as the preferred noun[14].

For the decoding analyses (Figure 3B,C), we selected 215 of the 252 untuned, noun-responsive cells (307 noun-responsive neurons minus the 53 noun-selective units, Table S2) for which we had more than four correct trials in every condition and which did not come from the first session with a different number of words.

Latency analysis

To measure the latency of the hippocampal responses elicited by nouns and pronouns, we compared the activity evoked by $Noun_{Pref}$ and $Noun_{NonPref}$ (or $Pronoun_{Ref}$ and $Pronoun_{NonRef}$) in a time-window from 0 to 1000ms, with a paired t-test in 99 time-bins of 10ms duration, after baseline correction based on the activity before the presentation of the noun (pronoun). The latency was estimated as the first of ten successive time-bins in which the response evoked by $Noun_{Pref}$ ($Pronoun_{Ref}$) was significantly higher than the response to the other nouns (pronouns). We used bootstrapping to determine the 95%-confidence interval for the latency (Fig. 2F,G), randomly selecting 53 noun-selective cells with replacement per bootstrap iteration ($N$ = 10,000).

Decoder analysis

We used a population decoding approach to examine information about nouns and pronouns at the population level. We trained decoders to discriminate between responses evoked by preferred and non-preferred nouns. For each iteration of the decoder, we selected 14 trials in which $Noun_{Pref}$ was present at the first location and 14 trials in which $Noun_{Pref}$ was present at the second location randomly by permutation for each unit, and held back two trials of each type for cross-validation. The analysis was based on the mean response to the nouns in a window from 200-700ms after word onset yielding a 56 (trials) x 50 (noun-selective cells) matrix (we excluded two cells from a session with a different number of words and one cell with too few correct trials to balance the decoder). We trained a linear support-vector machine (SVM) using the Matlab fitcsvm.m function (‘box constraint’ hyperparameter set to 1). We tested the output of the decoder elicited by the nouns on the held-back trials in sliding windows of 500ms duration, starting 500ms before the presentation of the first noun, and shifting the window by 100ms until the end of the second sentence (7000ms), yielding a total of 96 windows. To evaluate the performance of the decoder, we split the trials into three groups for both sentences. For the first sentence, we distinguished between trials with $Noun_{Pref}$ (i) at the first position, (ii) the second position and (iii) not present. For the second sentence, we distinguished between trials (i) in which the pronoun referred to $Noun_{Pref}$, (ii) with $Noun_{Pref}$ in sentence one but not referred to by the pronoun ($Pronoun_{PresNotRef}$) and (iii) with $Noun_{Pref}$ not present in sentence one ($Noun_{Absent}$; Fig. 3A and Fig S2B for testing on the $Pronoun_{AbsentSame}$ condition). We trained 96 decoders with different random samples of trials and tested them on the four held-out trials, yielding a total of 384 labels per time-window. The mean label was then taken as the output.

The decoder output represents the fraction of trials in which the classifier indicated the presence of $Noun_{Pref}$ in the time-window of interest. In Fig. 3A, for example, the decoder output reaches a level of 0.9. Hence, the decoder judged that the noun was present in this time-window on ~90% of individual test trials.

To assess the reliability of the classifiers at the population level, we generated 1000 surrogate populations of cells by selecting 50 cells from the pre-selected cells with replacement and determined the standard deviation across the 1000 surrogates. To determine the significance of differences between conditions, we compared decoder labels across time windows using a bootstrapped variant of a z-test. For each time window we calculated a z-statistic by dividing the difference between the classifier output for each of the two conditions to be tested by the standard deviation of this difference, across the 1000 surrogate populations. We determined a p-value based on a z-distribution (two-sided test). Because the test was performed for each of 96 time windows, we controlled for multiple comparisons using the false discovery rate correction[52].

To test the time generalisability of the decoding approach, we created a cross-decoder which was trained and tested on every combination of time-bins from the first and second sentence, in the same way as described above (Fig. S2C). For each time-bin we took the difference in the decoder label on trials when the preferred noun was present in the sentence from when the preferred noun was absent. We calculated the significance of the difference for each bin using the z-test described above. To correct for multiple comparisons, we clustered significant ($P < 0.05$, z-test) neighbouring time-bins. Clusters were considered to be significant if they passed a cluster-size threshold of greater than six time-bins, this threshold produces a FWE rate of 5% for independent samples.

We also tested decoders on the ambiguous trials, separately on trials on which the participant indicated that the pronoun referred to $Noun_{Pref}$ ($Chose_{Pref}$) or $Noun_{NonPref}$ ($Chose_{NotPref}$) (Fig. 3D). Units that preferred the unique gender could not be included in these decoders as $Noun_{Pref}$ was not present on ambiguous trials and 38 units remained for the analysis. The decoders were trained on the nouns of non-ambiguous trials and tested on ambiguous trials so that training and test trials were independent. We therefore did not need to hold out trials from the training set, resulting in more trials (32 noun present and 32 noun absent trials). The number of trials in the training set with the noun in the $1^{st}$ and $2^{nd}$ position in the first sentence was the same.

To examine the contribution of different cell types to the pronoun response, we trained decoders with only concept cells, NCNS cells or untuned cells and tested them on the responses evoked by $Pronoun_{Ref}$ or $Pronoun_{PrefNotRef}$ (time window from 200ms-700ms) (Fig. 3C). We also examined how the decoder output depended on the number of cells (Fig. 3C). We selected cells with replacement from different groups of neurons. To examine whether concept cells contain unique information we first selected from the concept cell class (n = 19), once all concept cells were included, we added cells from the NCNS class. As there were more NCNS cells ($N = 31$) than concept cells we selected a maximum of 19 NCNS cells from the population to allow a fair comparison of decoder performance between these groups. Finally, we selected cells from the untuned cells ($N = 215$). We also used a version in which we first added NCNS cells, then concept cells, and then untuned cells and adding cells in a random order (Fig. 3C).

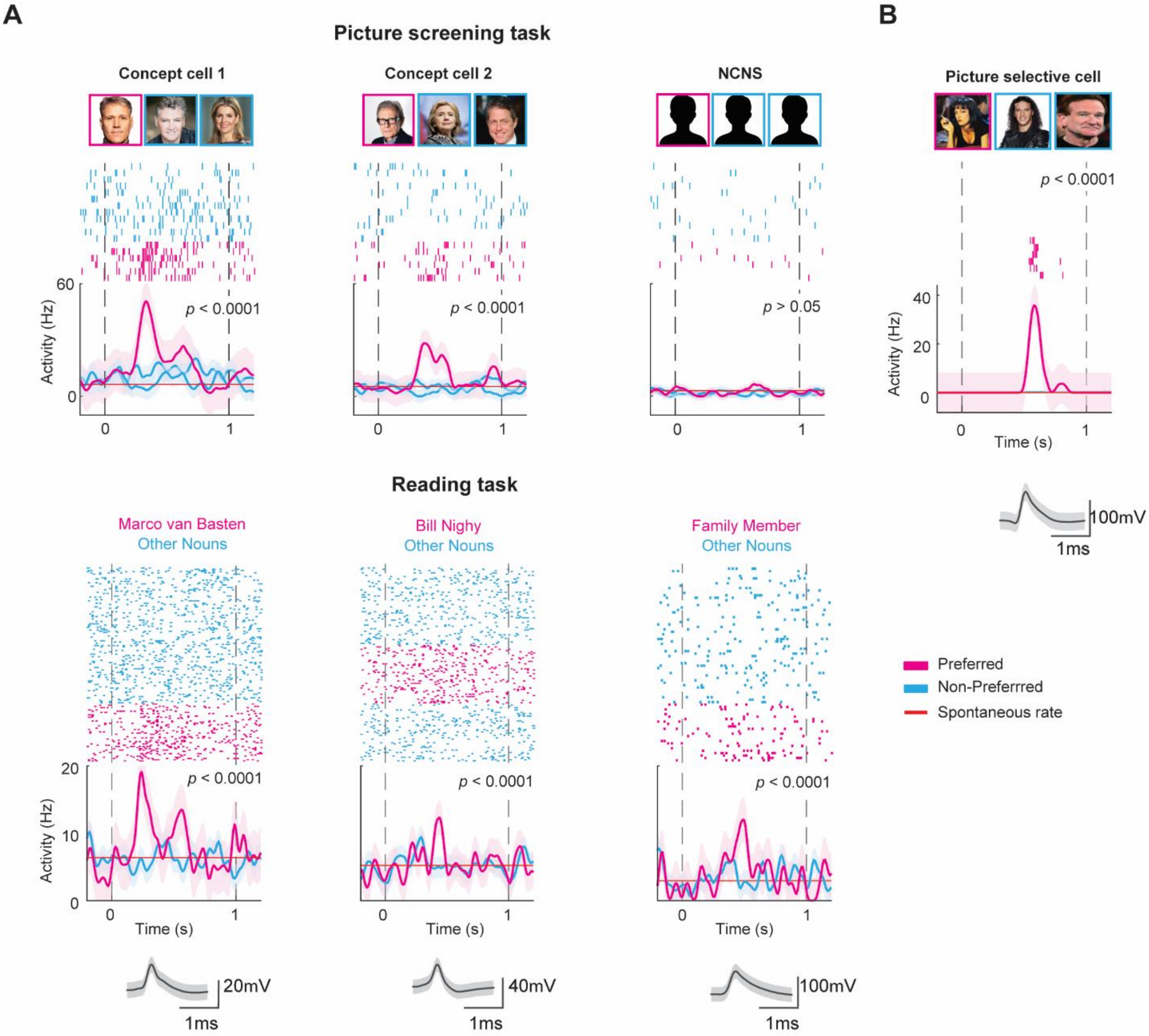


**Fig. S1. Example units.** **(A)** Two concept cells and one NCNS cell are shown with their average waveshape, their response to nouns during the reading task and to pictures during the screening task. Response to the preferred noun and picture are shown as a pink trace and the response to the non-preferred nouns and pictures as a blue trace. The p-values indicate the result of the permutation-based Poisson ANOVA. **(B)** One cell showing a selective response to a picture during the screening task. This cell did not exhibit noun selectivity during the reading task.

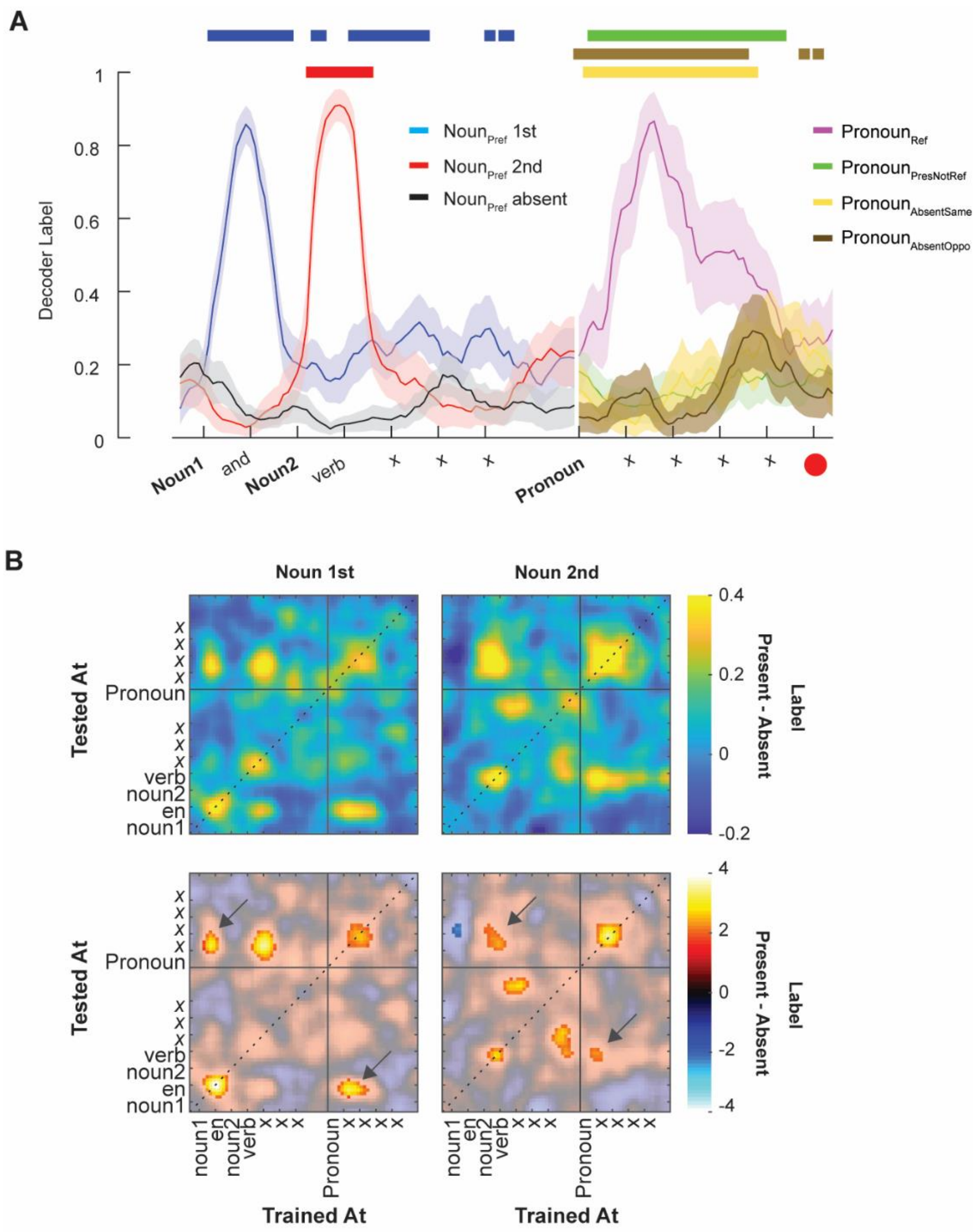


**Fig. S2. Decoder analyses**. **(A)** Decoder output for 45 neurons in which we could determine the response to $\text{Pronoun}_{\text{AbsentSame}}$. Filled bars, significant clusters of time windows ($P < 0.05$; see Methods). **(B) Left,** Output of the cross-decoder trained and tested on trials with $\text{Noun}_{\text{Pref}}$ at the first location. The colour-scale indicates the difference in decoder label between trials with the preferred noun and trials in which the preferred noun was absent in the first sentence. For trials with the preferred noun, we selected all the trials in which pronoun of the second sentence referred to it. Dashed diagonal, decoders trained and tested in the same time window. Note that the output of the decoder is a probability (i.e. a number between 0 and 1) but the difference between two conditions can become negative. **Right,** Output of the cross decoder trained and tested on trials with $\text{Noun}_{\text{Pref}}$ at the second location. **Top panels,** Output expressed as average label. **Bottom,** Z-scores. A cluster-based analysis shows significant clusters in the bottom two panels. Significant clusters are shown in bright colours.

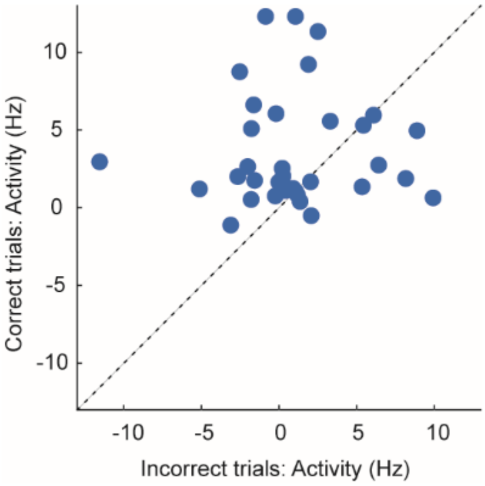


**Fig. S3. Relation between the hippocampal activity and the participants' answers.** We compared the activity elicited by pronoun referring to the preferred noun (Pronoun$_{Ref}$) between correct trials (y-axis) and trials on which the participant reported to wrong noun as subject of the second sentence (x-axis). We included 35 neurons with enough error trials. Most points were above the diagonal, which indicates that the hippocampal response was stronger on correct trials (two-tailed t-test, $t$ = -2.31, $p$ = 0.008).

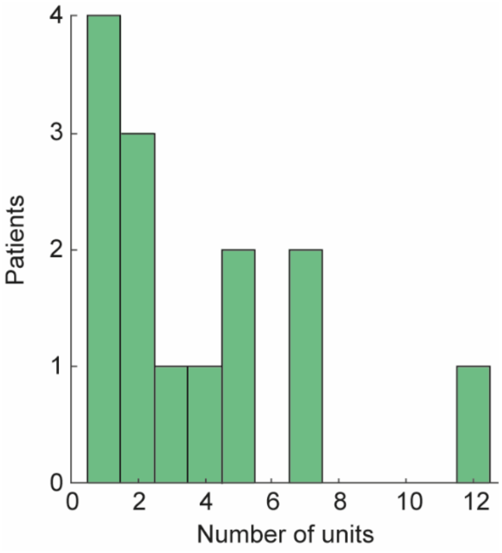


**Fig. S4. Number of cells per participant.** Number of cells that were recorded in each of the patients. One patient contributed 12 neurons, the others fewer.

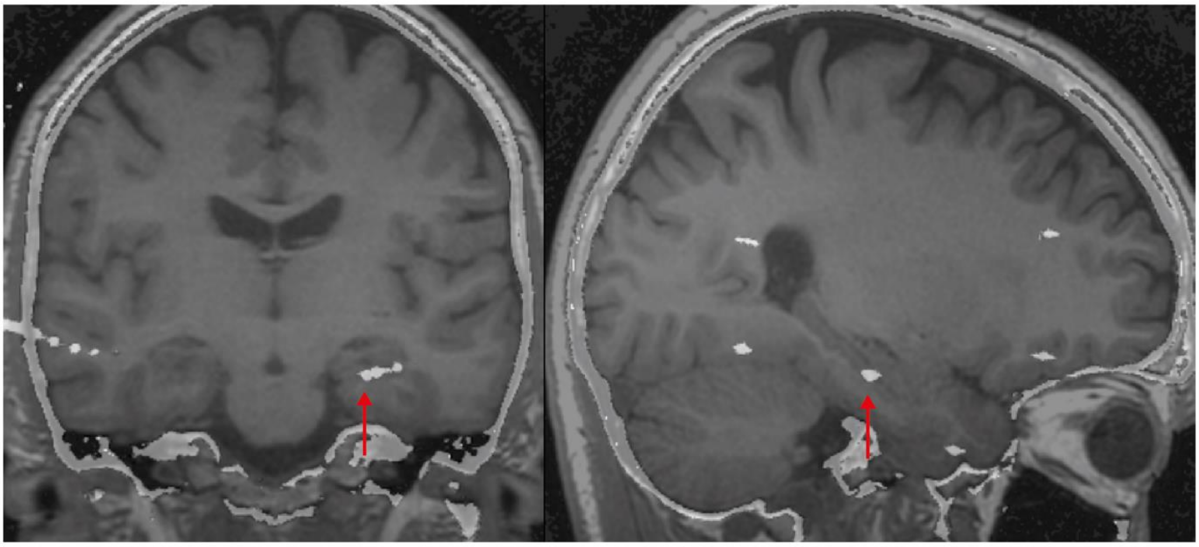

**Fig. S5. Position of the electrodes in the hippocampus.** Example post-implantation CT scan, in which the macro-contacts are visible, co-registered to the pre-implantation T1-weighted structural MRI scan. We confirmed that the microwires were in the hippocampus based on the visible macro-contacts (red arrow) and identified the end of the depth electrode from which the microwires extended.

| **Contexts** | First sentences: | Second sentences: |
|---|---|---|
| Dutch | $Noun_1$ en $noun_2$ liepen de bar binnen | *Pronoun* zat aan de tafel |
| | $Noun_1$ en $noun_2$ zaten in het park | *Pronoun* zette een zonnebril op |
| | $Noun_1$ en $noun_2$ keken naar de TV | *Pronoun* veranderde opeens de zender |
| | $Noun_1$ en $noun_2$ waren aan het eten | *Pronoun* opende een fles wijn |

**Table S1. Sentences for Dutch patients.**

| | Total in hippocampus | Mean per patient (N = 22) | Mean per session (N = 49) |
|---|---|---|---|
| All single and multi-units | 529 | 24.0 | 10.8 |
| Picture selective cells | 79 (15%) | 3.6 | 1.6 |
| Noun-responsive cells | 307 (56%) | 14.0 | 6.3 |
| Noun-selective cells | 53 (10%) | 2.4 | 1.1 |
| Concept cells | 19 (3.6%) | 0.9 | 0.4 |

**Table S2. Units recorded in the hippocampus.** A total of 529 single and multi-units were included. We recorded from 79 cells that were selective for pictures (tests are described in Methods). There were 307 units that responded to nouns and (increase in firing rate > 0.5 Hz) and we focused our analysis on 53 the noun-selective cells. Concept cells (N=19) exhibited picture selectivity and noun-selectivity for the same concept. Numbers between brackets are the percentage of the entire population.

| Patient ID | Reading sessions (*N* = 49) | sEEG probes in the hippocampus (*N* = 49) | Hippocampal units (*N* = 529) | Noun-responsive (*N* = 307) [in decoder] | Picture selective (*N*=79) | Noun-selective (*N* = 53) |
|---|---|---|---|---|---|---|
| *10* | 4 | 3 | 16 | 12 [0,0] | 4 | 3 [excl. in decod.] |
| *11* | 5 | 3 | 45 | 28 [0,0] | 8 | 0 |
| *21* | 1 | 1 | 2 | 2 [0,0] | 0 | 0 |
| *22* | 3 | 2 | 31 | 18 [18,16] | 7 | 2 |
| *23* | 2 | 2 | 25 | 12 [12,10] | 0 | 2 |
| *24* | 1 | 2 | 13 | 3 [3,3] | 0 | 0 |
| *25* | 2 | 2 | 7 | 7 [7,5] | 0 | 2 |
| *26* | 1 | 1 | 1 | 1 [1,1] | 0 | 0 |
| *29* | 3 | 2 | 49 | 40 [40,33] | 4 | 7 |
| *30* | 1 | 2 | 7 | 6 [6,5] | 1 | 1 |
| *33* | 1 | 2 | 13 | 3 [3,2] | 4 | 1 |
| *34* | 2 | 2 | 36 | 25 [25,20] | 5 | 5 |
| *37* | 3 | 2 | 40 | 21 [21,20] | 3 | 1 |
| *38* | 5 | 2 | 32 | 13 [13,9] | 8 | 4 |
| *39* | 3 | 2 | 54 | 35 [35,30] | 14 | 5 |
| *40* | 2 | 1 | 4 | 1 [1,1] | 0 | 0 |
| *41* | 2 | 2 | 11 | 8 [8,8] | 1 | 0 |
| *102* | 2 | 4 | 64 | 28 [28,21] | 4 | 7 |
| *104* | 1 | 4 | 4 | 4 [4,4] | 0 | 0 |
| *106* | 1 | 2 | 11 | 3 [3,3] | 1 | 0 |
| *207* | 3 | 4 | 54 | 32 [32,20] | 10 | 12 |
| *208* | 1 | 2 | 10 | 5 [5,4] | 5 | 1 |

**Table S3. Units, sessions and electrodes per patient.** The total number of noun-responsive units was 307. The first and second number between brackets in this column refer to the total number of cells and the number of untuned cells that could be included in the decoder, respectively. Three noun-selective cells of participant 10 were excluded from the decoder because the sentences contained a different number of words.
